# Genome-wide identification of alternative splicing events that regulate protein transport across the secretory pathway

**DOI:** 10.1101/459263

**Authors:** Alexander Neumann, Magdalena Schindler, Didrik Olofsson, Ilka Wilhelmi, Annette Schürmann, Florian Heyd

## Abstract

Alternative splicing (AS) strongly increases proteome diversity and functionality in eukaryotic cells. Protein secretion is a tightly-controlled process, especially in a tissue-specific and differentiation-dependent manner. While previous work has focussed on transcriptional and posttranslational regulatory mechanisms, the impact of AS on the secretory pathway remains largely unexplored. Here we integrate a published screen for modulators of protein transport and RNA-Seq analyses to identify over 200 AS events as secretion regulators. We confirm that splicing events along all stages of the secretory pathway regulate the efficiency of protein transport using Morpholinos and CRISPR/Cas9. We furthermore show that these events are highly tissue-specific and adapt the secretory pathway during T-cell activation and adipocyte differentiation. Our data substantially advance the understanding of AS functionality, add a new regulatory layer to a fundamental cell biological process and provide a resource of alternative isoforms that control the secretory pathway.

## Introduction

After biosynthesis in the endoplasmic reticulum (ER), proteins are transported to their destination via the secretory pathway. Transport processes are highly flexible and adapt to new requirements for instance during differentiation or after activation (Farhan and Rabouille, 2011; McCaughey and Stephens, 2018). This adaptation is achieved either by differential gene expression (Coutinho et al., 2004; Dunne et al., 2002; Schotman et al., 2009), by changes in membrane morphology and dynamics (Farhan et al., 2008; Forster et al., 2006; Guo and Linstedt, 2006) or by altered activity of kinases and phosphatases (Farhan et al., 2010). Alternative splicing has been shown to react highly dynamic to various external stimuli (Heyd and Lynch, 2011; Preußner et al., 2017) and would thus be perfectly suited to control protein secretion in response to changing cellular environments. Indeed, we have previously described how AS of SEC16A exon 29 increases the efficiency of the early secretory pathway (Wilhelmi et al., 2016) but this remains one isolated example. Here, we globally identify AS events that modulate the secretory pathway in a tissue-specific and activation‐ and differentiation-dependent manner.

## Results and discussion

To identify regulators of the secretory pathway, several RNAi screens have been performed (Farhan, 2015). One of these screens used the VSVG reporter to monitor protein transport upon knock-down (KD) of 22,000 human genes (Simpson et al., 2012). In this screen, four RNA-binding proteins (RBPs) that regulate AS and whose KD led to a strong inhibition of secretion were identified (Figure 1 – Source Data 1): HNRNPA1, PTBP1, RBM27 and SRSF1 (Figure 1A). As these RBPs have no known direct function in protein secretion, we considered an indirect effect, through controlling AS of components of the secretory pathway, which then changes cargo flux (Figure 1B). We therefore used KD RNA-Seq datasets of these RBPs to determine their splicing targets. We propose that the impact of these RBPs on the secretory pathway is mediated by a network of common AS events and thus generated an overlap of splicing targets that were differential in the single KDs. We defined those that were regulated by at least three of the four RBPs as potential secretion modulators, henceforth called secretion events (Figure 1C). Meta-analysis of these events, which are mostly cassette exons, indicates that they have a modulatory effect on protein function instead of regulating total gene expression, as they lie almost exclusively within the coding sequence and consist of less NMD (nonsense-mediated decay)-inducing exons when compared to the skipped exons that are regulated by any of these RBPs (RBP background, Figure 1 – figure supplement 1). When performing gene ontology (GO) analysis, only three terms for biological processes were found significantly enriched, all of which fit the hypothesis that the secretion events play a role in membrane trafficking (Figure 1D). We then grouped the AS-controlled proteins by function (Figure 1E). As expected, a large cluster is directly connected to the secretory pathway or to cytoskeletal organizers that can provide scaffolding functions for vesicle transport. Additionally, we find phosphoinositide homeostasis and ARF signalling, which are known to regulate vesicular trafficking (Ooms et al., 2009) and proteins involved in kinase and phosphatase signalling, which can post-translationally control trafficking proteins (Farhan et al., 2010). Apart from these targets, there are several groups that may have an indirect effect on secretion: transcriptional, post-transcriptional and translational modulators can change gene and protein expression of various targets, which can be further controlled by degradation pathways. Mitochondrial proteins could impact on mitochondria-related traffic. Finally, there are several chaperones, a group of proteins which have been shown to influence protein secretion as well (Roth et al., 2012). Closer inspection of the proteins known to be direct part of the trafficking machinery revealed that they are localized in all compartments along the secretory pathway (Figure 1E).

**Figure 1:**
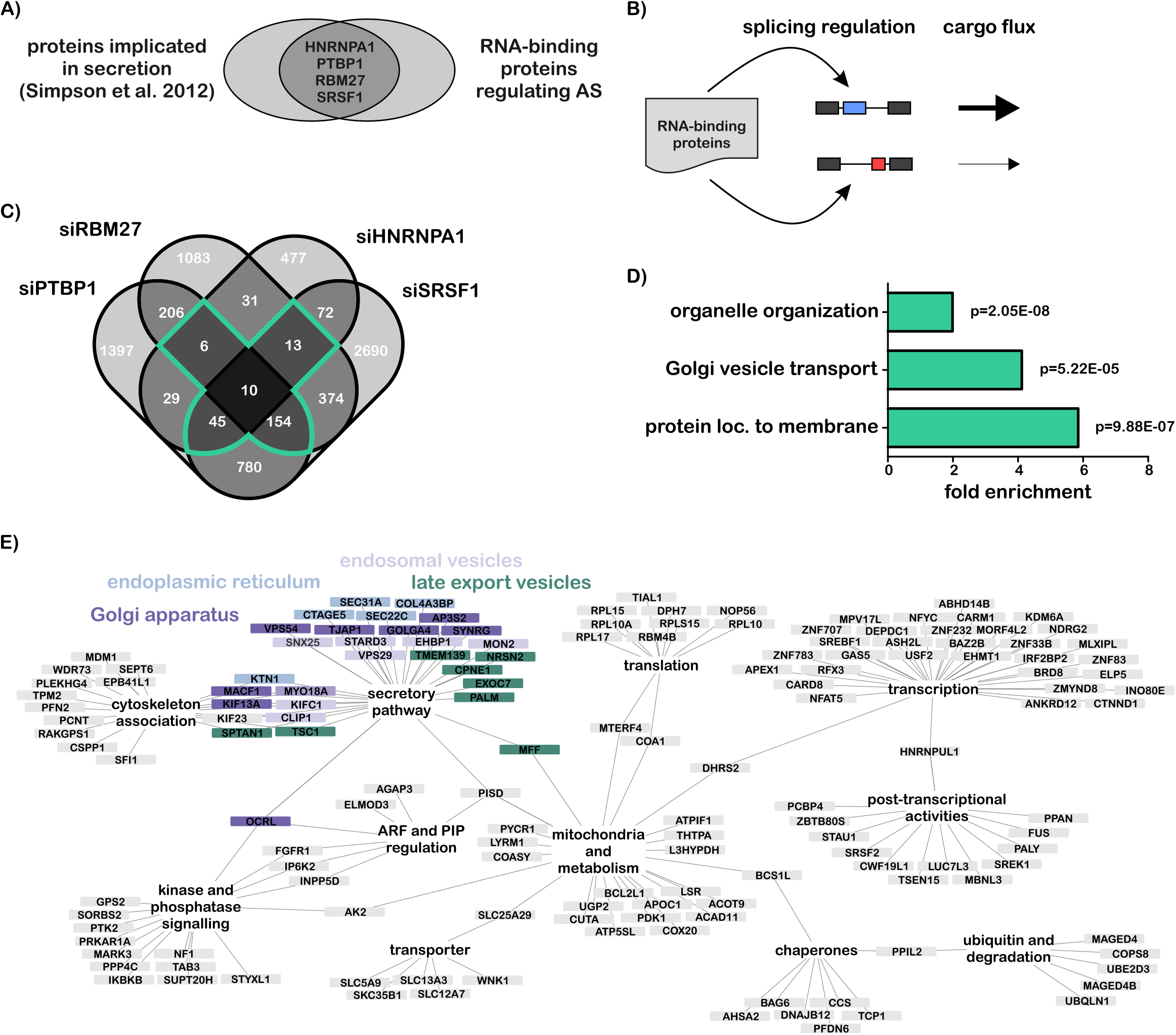
Genome-wide identification of secretion-modulating alternative splicing events. **A)** Venn diagram of proteins implicated in secretion phenotypes upon KD and alternative splicing (AS)-regulatory RNA binding proteins. **B)** Model of how RBPs indirectly regulate cargo flux by AS. **C)** Venn diagram of RBP KD RNA-Seq AS analyses. Events that are differentially spliced in at least three KDs are candidates for secretion regulators (marked in green, termed secretion events). **D)** Biological process GO terms of secretion event genes (all significantly enriched terms shown) with all genes that change splicing in any of the KDs as background (RBP background). **E)** Network of the secretion events describing their function according to RefSeq gene annotations. Proteins of the secretory pathway are color-coded based on their localization.

To validate the functionality of the secretion events we employed the RUSH system (Boncompain et al., 2012) in which a GFP-GPI reporter is transported from the ER to the PM after addition of biotin. Upon arrival of the reporter at the PM, GFP will be located on the extracellular side and can be quantified by staining against GFP without permeabilizing the cell (Figure 2A). A time course experiment in HEK293T cells and the quantification of antibody-stained (PM-localized) GFP per cell is shown in Figure 2B. To verify that expression levels of HNRNPA1, PTBP1, RBM27 and SRSF1 had an effect on protein transport in our assay, we performed KDs of these proteins and of two further RBPs (MBNL3, SRSF6) as a control (Figure 2C). The RUSH assay was subsequently performed with GFP staining 1 h after biotin addition. While we did not observe a significant difference between control and either the MBNL3, SRSF6 or HNRNPA1 KDs, cells treated with siPTBP1, siRBM27 or siSRSF1 showed a substantial and highly significant decrease in GFP surface staining (Figure 2D, E). The lack of an influence of the HNRNPA1 KD (Simpson et al., 2012) may be due to the use of a different reporter in our study, which could point to a role in cargo selectivity. However, we also notice that HNRNPA1 has the least common splicing targets among the four RBPs, as the largest overlap in our analysis are the targets controlled by PTBP1, RBM27 and SRSF1 (Figure 1C). For further validations, we therefore focussed on events that were controlled by these three RBPs.

**Figure 2:**
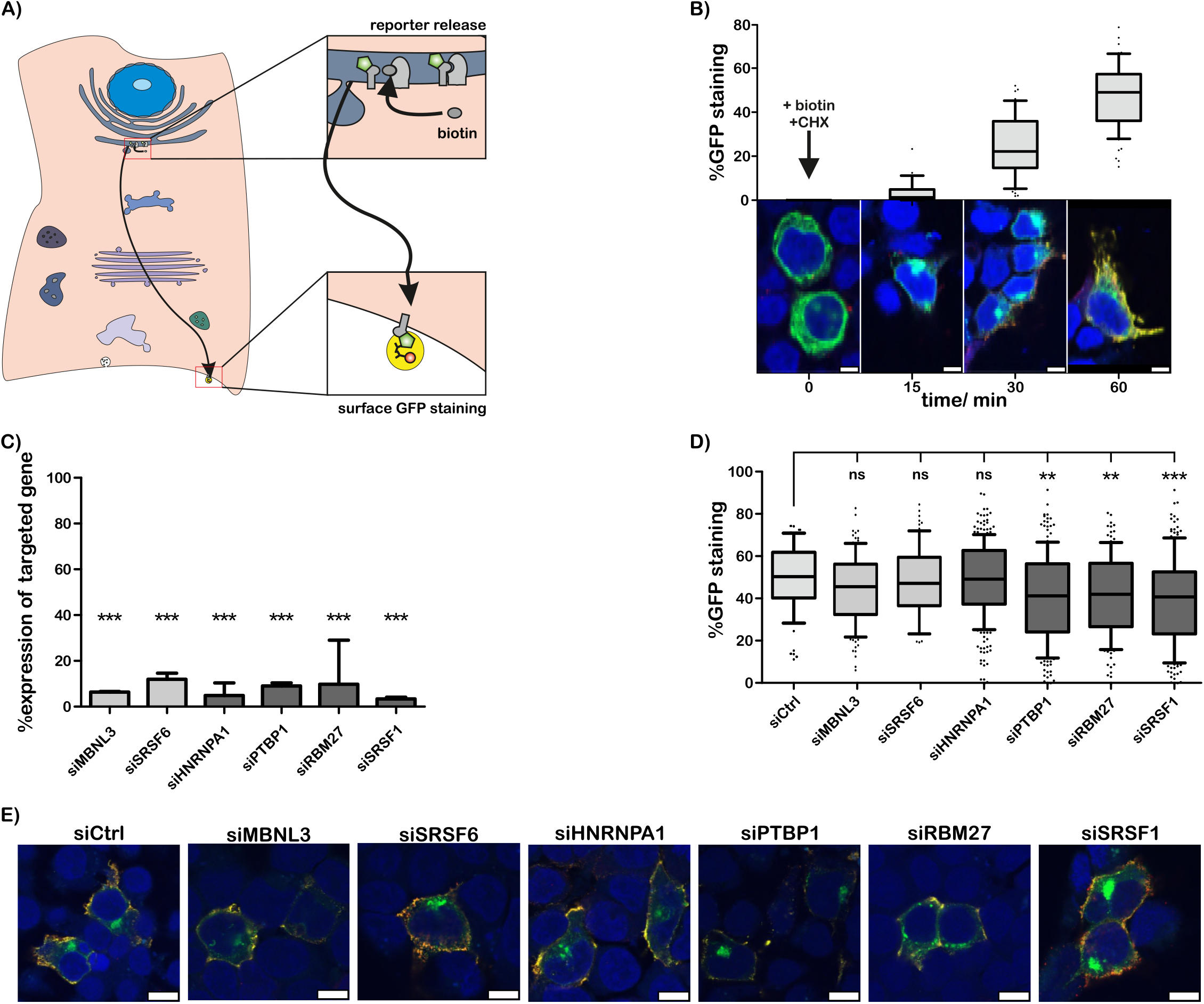
Validation of a secretion phenotype upon RBP KD. **A)** The RUSH system principle with extracellular GFP staining as readout for transport efficiency. **B)** Time course experiment with quantification of the RUSH assay. n>=21 cells, significances not shown. **C)** Validation of the RBP KD. Shown are %expression of each targeted gene, normalized to GAPDH and control siRNA. Shown are mean and SD, n>=3. **D)** Quantification of the RUSH assay in control and RBP KD cells. Whiskers show 10-90 percentiles, n>=68 cells. **E)** Representative images corresponding to d. ns p>=0.05, ** 0.01>p>=0.001, *** 0.001 >p, scale bar 10 μm.

To validate the influence of individual splicing events on secretion, we selected four targets that are directly involved in different parts of the secretory pathway (Figure 1E): for the early secretory pathway, we selected SEC31A, which is part of the outer coat of COPII vesicles (Gürkan et al., 2006) and SEC22C, which is a homolog of the SNARE protein Sec22 in yeast (Tang et al., 1998; Yamamoto et al., 2017). As examples for splicing events affecting Golgi and post-Golgi components of the secretory pathway, we selected OCRL, which acts at the *trans-Golgi* and later compartments as a phosphoinositide phosphatase (De Matteis et al., 2017) and EXOC7 (also known as EX070), which is part of the exocyst complex involved in targeting vesicles to the PM (He and Guo, 2009). We used splice-site blocking Morpholinos (MOs) to manipulate AS of these targets without altering the endogenous gene expression level (Figure 3A) and performed RUSH assays on these cells. We indeed observed a highly significant decrease in GFP reporter transported to the PM for all candidates (Figure 3B, C).

**Figure 3:**
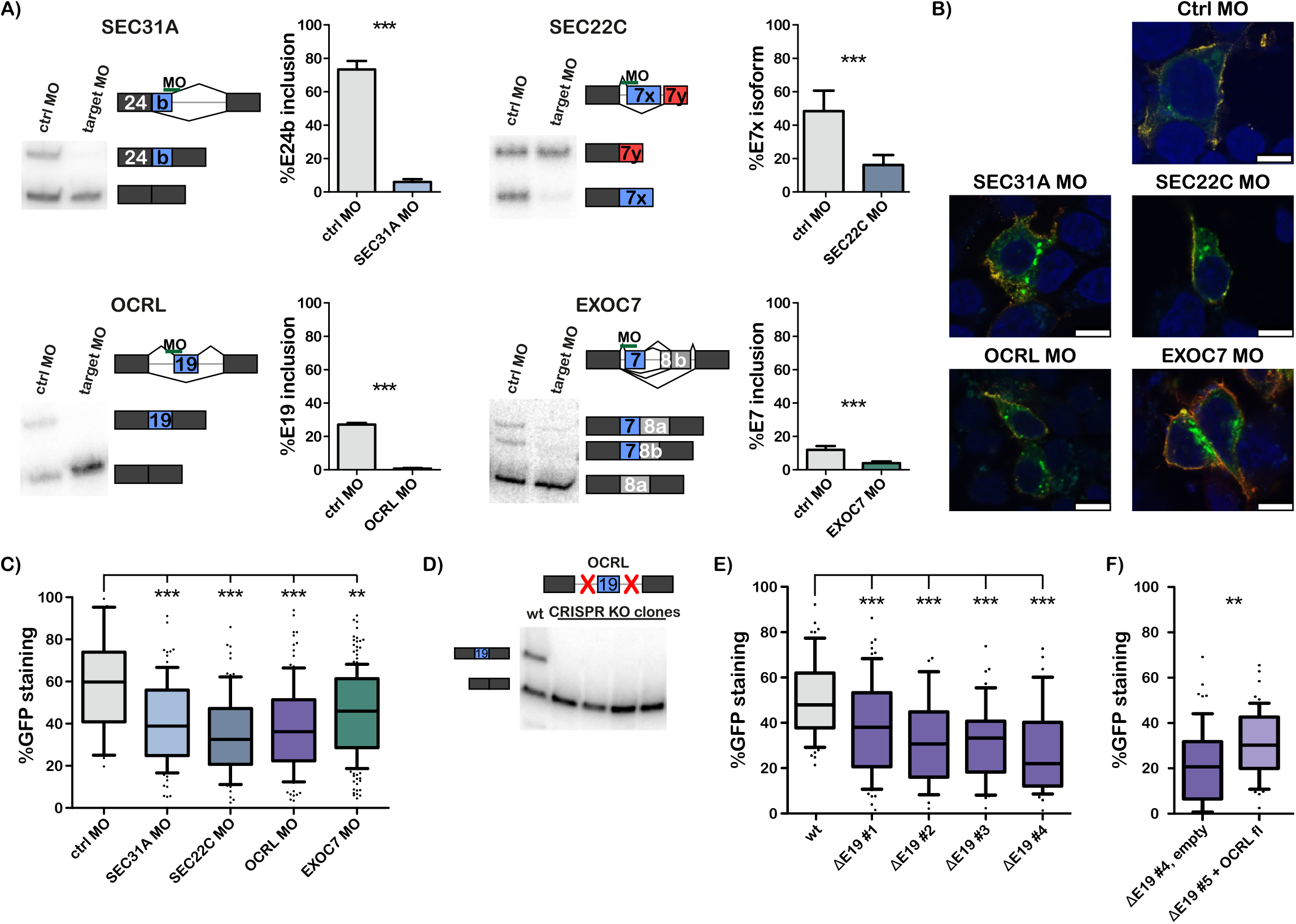
Validation of a secretion phenotype upon manipulation of AS secretion events. **A)** Representative radioactive RT-PCR and quantifications of cells treated for 48 hours with Morpholinos against SEC31A exon 24b, SEC22C exon 7x, OCRL exon 19 and EXOC7 exon 7, respectively. Shown are mean and SD, n>=3. **B)** Representative microscopy images of cells treated with control MOs and MOs against the targeted exons from a. Scale bar 10 μm. **C)** Quantification of the RUSH assay performed with cells pre-treated with MOs for 36 hours before transfection of the RUSH plasmid. n>=29 cells. **D)** Validation of CRISPR/Cas9-mediated knock-out of OCRL exon 19 for four independent cell lines on transcriptomic level using radioactive RT-PCR. **E)** Quantification of the RUSH assay performed with wild-type and OCRL exon 19 knock-out cells. n>=25 cells. **F)** RUSH assay quantification. Tested was OCRL exon 19 KO cell line #4 with or without OCRL full-length expression. n>=60 cells. Box plot whiskers show 10-90 percentiles, ns p>=0.05, ** 0.01>p>=0.001, *** 0.001 >p.

To independently validate the observed effect, we used CRISPR/Cas9 to generate isoform-specific knock-outs (KOs). We selected the alternative microexon 19 in OCRL, as it stands out both because the MO-induced splicing change from about 25% inclusion to complete exclusion led to a drastic transport defect and as this is achieved by exclusion of only 8 amino acids. We generated four independent OCRL exon 19 KO cell lines and validated them both on genomic and transcriptomic level (Figure 3 – figure supplement 1A, Figure 3D). Cell lines showed expression of only the exclusion isoform with the overall expression level remaining largely unchanged (Figure 3 – figure supplement 1B). We then performed the RUSH assay and again observed a highly significant reduction of surface-GFP for all clones in the same range as for the OCRL exon 19 MO-treated cells, further validating the MO approach (Figure 3E). In addition, when expressing OCRL full-length protein for a short period (Figure 3 – figure supplement 1C) in the KO cells, the defect in protein secretion was partially rescued (Figure 3F), while there was no effect observed in wild-type cells (Figure 3 – figure supplement 1D). These data together validate our bioinformatics approach, as all tested splicing events indeed control protein transport efficiency. This strongly increases the confidence in our group of secretion events, thereby adding a new regulatory layer to the secretory pathway and substantially increasing the number of alternative isoforms with known cellular functionality.

To address the physiological significance of the connection between AS and protein transport, we investigated whether the secretion events act in a tissue-, differentiation‐ or activation-dependent manner. To this end, we initially used RNA-Seq data from various human organs to calculate inclusion levels for the secretion and RBP background events. We indeed observed a tissue-specific usage for a larger proportion of the secretion events in comparison to the RBP background (Figure 4A, Figure 4 – figure supplement 1A). When calculating measures of variability between tissues, more secretion events were found differential in any two-way comparison (Figure 4B) and the strongest PSI (percent spliced in) difference of an event between tissue was larger (Figure 4C). This strongly points to a global role of AS in adapting the secretory pathway to tissue-specific requirements. Next, we turned to two cellular systems where cells with basal secretory requirements differentiate into a cell type with higher secretory load (Figure 4D): T-cell activation and differentiation of pre-adipocytes into adipocytes. We analysed differential AS using RNA-Seq datasets from primary human CD4^+^ T-cells and human SGBS adipocytes pre‐ and post-differentiation and found significant overlaps with the secretion events in both sets (Figure 4E, F) from which we validated three targets each (Figure 4 – figure supplement 1B, C). Of note, the vast majority of secretion-related AS events act independently of transcription, as about 80% of the corresponding genes show a fold-change smaller than 2 (Figure 4 – Source Data 1). We observed that while the T-cell overlap events are found in all parts of the secretory pathway, the adipocyte overlap events mainly locate in post-Golgi compartments, which is also reflected in GO term analysis (Figure 4 – figure supplement 1D). Additionally, we find that components of the COPII machinery are upregulated during adipocyte differentiation but not during T-cell activation (Figure 4G, Figure 4 – Source Data 1). This suggests that adipocytes use transcriptional regulation in the early steps of protein secretion and alternative splicing in post-Golgi compartments to adapt their secretory pathway, whereas T-cells rely on AS to increase secretion capacity during activation. This difference in the control of membrane trafficking may be explained by specific requirements of these cells after differentiation. Activated T-cells produce cytokines and cytotoxins at the ER and then transport them out of the cell, using the whole secretory pathway, which happens in a highly dynamic and temporally-controlled manner (Huang et al., 2013). Adipocytes produce and export adipokines and need to adapt their secretory machinery in this regard as well (Kuryszko et al., 2016), but this happens during a longer differentiation process that is more suitable for stable transcriptional changes. However, a fundamental task in mature adipocytes is the rapid shuttling of the glucose transporter GLUT4 from post-Golgi vesicles to the PM in response to insulin and the recycling of the receptor (Stöckli et al., 2011). A major and more dynamic adaptation is therefore required for the post-Golgi trafficking machinery, which is achieved by AS. These data together strongly argue for a global role of AS in controlling the efficiency of the secretory pathway in a tissue-specific manner, as well as in dynamic settings such as differentiation and activation (Figure 4 – figure supplement 1E).

**Figure 4:**
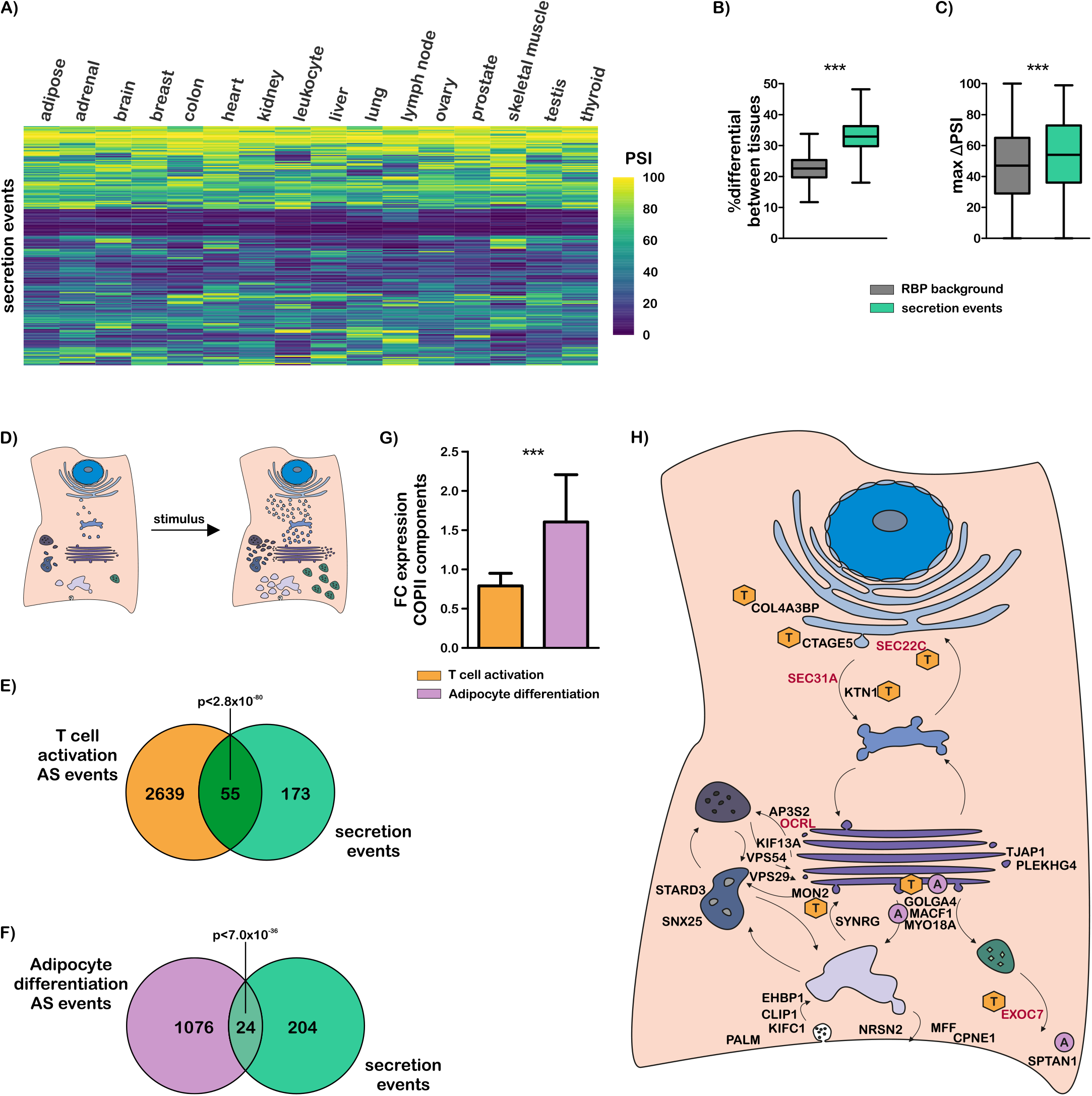
Tissue‐ and differentiation-specific usage of secretion events to adapt the secretory pathway. **A)** Tissue heatmap of secretion events based on RNA-seq data from the lllumina bodymap. PSI percent spliced in. **B)** For every possible two-way tissue comparison, the percent of differential events was calculated. Shown are the secretion events and the RBP background. **C)** Maximal PSI difference (ΔPSI) between any tissues per event for the secretion events and the RBP background. **D)** Model of a cell with low secretory potential that differentiates Into a cell with high secretory potential after a stimulus. **E, F)** Overlap of T-cell activation-specific (E) and adipocyte differentiation-specific (F) AS events with the secretion events, p values calculated via hypergeometric distribution. **G)** Gene expression analysis of COPII components from the RNA-Seq data of T cells (orange) and adipocytes (violet) pre‐ and post-dlfferentlatlon. FC fold change. **H)** Proposed model of splicing-regulated secretion modulators. Proteins of the secretion events linked to the secretory pathway are shown, with their position Indicating their function. Validated events are marked In red and differentiation-specific events with an orange T and violet A for T-cell activation‐ and adipocyte differentiation-specific, respectively. *** p<0.001. Whiskers show maximal and minimal value.

In summary, we combine data from a genome-wide screen for modulators of protein secretion with knock-down RNA-Seq datasets to discover over 200 AS events that, based on our validations using the RUSH-system, are high-confidence secretion regulators. Our analysis shows that AS is involved in regulating all stages of the secretory pathway (Figure 4H). We furthermore show that secretion-regulating AS events are used in diverse biological contexts to adapt the secretory pathway to tissue-specific or differentiation‐ and activation-dependent requirements (Figure 4H, Figure 4 – figure supplement 1E). These findings extend the analysis of individual splicing events with a role in membrane trafficking (Blue et al., 2018; Valladolid-Acebes et al., 2015; Wilhelmi et al., 2016) to a systems-wide level and add a new layer of complexity to the regulation of the secretory pathway. In addition, the dynamic nature of these splicing events in various biological contexts provides an important step towards understanding differentiation-specific control of protein secretion. Alternative splicing might help the cell to adapt the secretory pathway in an intermediate timeframe, supplementing the very fast modulations of kinases and phosphatases and the slower effects of transcription. Despite clear variations in cargo load and specificity in different cells and tissues, the molecular basis for these distinct adaptations remains largely enigmatic. Our data provide evidence that cell-type-specific adaption of protein secretion is, at least partially, controlled by a network of splicing changes. Our finding that several RBPs are involved in controlling this process is consistent with splicing decisions being under combinatorial control of several or many *cis*‐ and *trans*-acting factors and explains how different cell types can adapt their splicing patterns to the respective requirements. The expression and activity of these RBPs can be individually controlled to result in a variety of activities that is tailored to the secretory requirement of the respective cell type.

## Materials and methods

### Accession numbers, RNA-Seq analysis and post-analysis

Accession numbers and all results from analyses are listed in Figure 1 – Source Data 1 and Figure 4 – Source Data 1. RNA-Seq analyses were essentially performed as previously described (Herdt et al., 2017). In short, reads were mapped to the hg38 human genome using STAR version 2.5.3a (Dobin et al., 2013). For alternative splicing analyses, mixture of Bayesian inference model (MISO) (Katz et al., 2010) version 0.5.3 with a custom-made annotation was used. Differential events are defined by a minimal ΔPSI of 10% and a Bayes factor greater 5. Replicates were merged for analyses after mapping. Gene expression analysis was performed using DESeq2 (Anders et al., 2012). Further analyses were performed using custom Python scripts. GO term analyses were performed using the PANTHER classification system version 13.1 (https://pantherdb.org) (Mi et al., 2013). Network maps were generated using Cytoscape version 3.6.1 (Montojo et al., 2010).

### Cell culture, transfections and genome-editing

HEK293T cells were cultivated in DMEM high glucose (Biowest, Nuaillé, France) containing 10% FCS (Biochrom, Berlin, Germany) and 1% Penicillin/ Streptomycin (Biowest) at 37 °C and 5% CO_2_. Plasmid transfections were performed using RotiFect (Carl Roth, Karlsruhe, Germany) following manufacturers’ instructions. Morpholinos were obtained from Gene Tools. They were transfected using Endo-Porter following manufacturers’ instructions. For KDs, a pool of four siRNAs (Dharmacon, Lafayette, Colorado, US) was used at final concentration of 10 nM. They were transfected using HiPerFect (Qiagen, Venlo, Netherlands) according to the manual. Genome-editing using CRISPR/Cas9 was performed as previously described(Wilhelmi et al., 2016). All sequences for Morpholinos, siRNAs, guide sequences and genotyping primers are listed in the Source Data files of Figures 2-4.

Human SGBS cells were cultivated and differentiated as described previously (Fischer-Posovszky et al., 2008; Wabitsch et al., 2001). In brief, cells were seeded in DMEM/F12 containing 10% FCS at a density of 4×10^4^ cells per 6-well for 3 days to reach confluency. Differentiation was started by application of a specific cocktail (2 μmol/l rosiglitazone, 25 nmol/l dexamethasone, 0.5 mmol/l methyliso-buthylxantine, 0.1 μmol/l cortisol, 0.01 mg/ml transferrin, 0.2 nmol/l triiodotyronin, and 20 nmol/l human insulin) in DMEM/F12 without serum and albumin. Medium was changed every 4 days (DMEM/F12 0.1 μmol/l cortisol, 0.01 mg/ml transferrin, 0.2 nmol/l triiodotyronin, and 20 nmol/l human insulin). Cells were harvested in Qiazol 3 days after seeding (pre-adipocytes) and 14 days post differentiation (adipocytes) for RNA extraction using the RNeasy MinElute Kit (Qiagen) according to manufacturers’ instructions.

### Constructs

The RUSH plasmid was generously provided by Franck Perez and consists of a KDEL ER hook and an EGFP-GPI reporter (Addgene #65294). The OCRL full length expression construct was purchased from Addgene (#22207).

### RNA extraction, RT-PCR, RT-qPCR and PCR

RNAs from primary human T-cells were prepared as described(Michel et al., 2014). RNA extraction, radioactive RT-PCR and Phosphor imager quantification were performed as previously described(Wilhelmi et al., 2016). RNA was extracted using RNATri (Bio&Sell, Feucht, Germany). For reverse transcription, 1 μg RNA was used with a gene-specific reverse primer. Low-cycle PCRs were performed with a radioactively-labelled forward primer and products were separated by denaturing PAGE. Imaging was done using a Phosphoimager and gels quantified using ImageQuantTL version 8.1 (GE, Boston, Massachusetts, US). qPCRs were performed in technical duplicates in a 96-well format using Absolute QPCR SYBR Green Mix (Thermo Fisher, Waltham, Massachusetts, US) on a Stratagene (San Diego, California, US) Mx3000P machine. Mean values of the technical duplicates were used and expression normalized to GAPDH. Non-radioactive genomic PCR products were separated on an agarose gel. See figure legends for number of independent biological replicates. Primer sequences are provided in the Source Data files of Figures 2-4.

### RUSH assay and staining

The day before transfection, cells were seeded on precision coverslips (0.17 mm thickness, Sigma, Kawasaki, Japan) that had previously been coated with poly-L-Lysine (Sigma) for 1 hour. For the normal RUSH assay, 1.0 × 10^5^ cells were seeded. 24 hours post-seeding, cells were transfected with the RUSH plasmid. 16 hours post-transfection, protein synthesis was stopped by addition of cycloheximide (10 μg/ml final concentration, Carl Roth) and D-biotin (40 final concentration, Sigma) was added to release the reporter. The assay was stopped, usually after one hour if not indicated otherwise, by washing with ice-cold PBS (Biowest) and fixation of cells using 4% formaldehyde (Carl Roth) solution in PBS for 10 minutes. After 1 hour of blocking using 5% goat serum (Sigma) in PBS, primary rabbit anti-GFP antibody (1:200, Invitrogen, Carlsbad, California, US) was applied for 2 hours at room temperature. After washing, the secondary donkey anti-rabbit AlexaFluor594 antibody (1:200, Invitrogen) was applied for one hour. Cover slips were mounted on microscopy slides (Roth) with ProLong Gold antifade mountant with DAPI (Thermo Fisher). Slides were stored at room temperature for 24 hours and either directly imaged or stored at 4 °C until imaging. When the RUSH assay was combined with either Morpholino or siRNA treatment, only 0.5 × 10^5^ cells were seeded to account for the additional day of treatment. MOs or siRNAs were transfected 32 hours before the RUSH plasmid. For the OCRL full length rescue experiments, the OCRL full length construct was transfected together with the RUSH plasmid.

### Microscopy and analysis

Fluorescence microscopy was performed on a Leica SP8 confocal microscope. Image analysis was performed using custom Python scripts. To calculate percent stained GFP, single cells were initially defined by user input. Pixels were defined GFP-positive (green value) or GFP staining-positive (red value) if the respective intensity value was above a certain threshold (75 for green, 65 for red fluorescence). %GFP staining was calculated as the number of double-positive pixels divided by the number of green-positive pixels multiplied by 100.

### Immunoblotting

Whole cell lysates were prepared in RIPA buffer (10 mM TrisHCL pH 8.0, 2 mM EDTA, 100 mM NaCl, 1% NP-40, 5 mg/mL sodium deoxycholate, with protease inhibitors). SDS-PAGE and western blotting followed standard procedures. Antibodies used: mouse monoclonal anti-HA-probe (F-7, 1:1000, Santa Cruz Biotechnology, Dallas, Texas, US), mouse anti-Gapdh (1:1000, antibodies online, Aachen, Germany) and horse anti-mouse IgG, HRP-linked (1:5000, Cell Signaling, Danvers, Massachusetts, US)

### Statistics

If not mentioned otherwise, either one-sample or unpaired T tests were used to calculate p values. P values are indicated by asterisks and explained in the legend of each figure. N numbers in the legends represent biological replicates. P value calculations were only performed if at least three biological replicate values were obtained.

## Acknowledgements

The authors thank the HPC Service of ZEDAT, Freie Universität Berlin, for computing time. We thank Franck Perez for generously providing the RUSH plasmid and Rainer Pepperkok for sharing unpublished information of the genome-wide screen performed in his lab. We also thank members of the Heyd lab for discussions and comments on the manuscript. This work was supported by the DFG SFB958/A21 to F.H.

## Source Data files

Figure 1 – Source Data 1: RNA-seq alternative splicing analysis and meta-analysis.

Figure 2 – Source Data 1: Raw data regarding Figure 2.

Figure 3 – Source Data 1: Raw data regarding Figure 3.

Figure 4 – Source Data 1: RNA-seq alternative splicing and gene expression analyses and meta analyses.

---

**Figure 1 - figure supplement 1:**
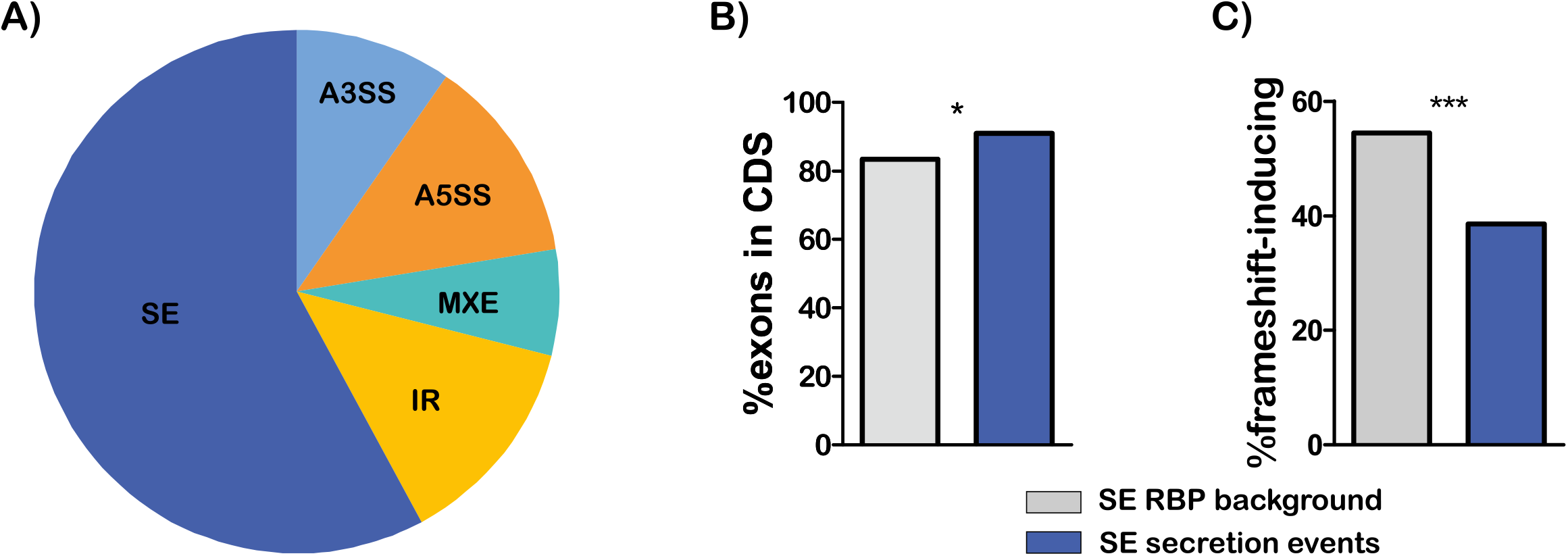
Meta-analysis of the secretion events. **A)** Distribution of event types for the secretion events into alternative 3′ and 5′ splice site (A3SS, A5SS), mutually exclusive exons (MXE), retained Introns (IR) and skipped exons (SE). **B, C)** Meta-analysis of skipped exon secretion events versus all skipped exons that are regulated by any of the four RBPs (RBP background). Exons In the coding sequence (CDS, B) and proportion of exons with a length not divisible by three, which are therefore frameshlft-lnduclng and potential nonsense-mediated decay targets (C). * 0.01<=p<0.05, *** p<0.001, Chi square test.

**Figure 3 - figure supplement 1:**
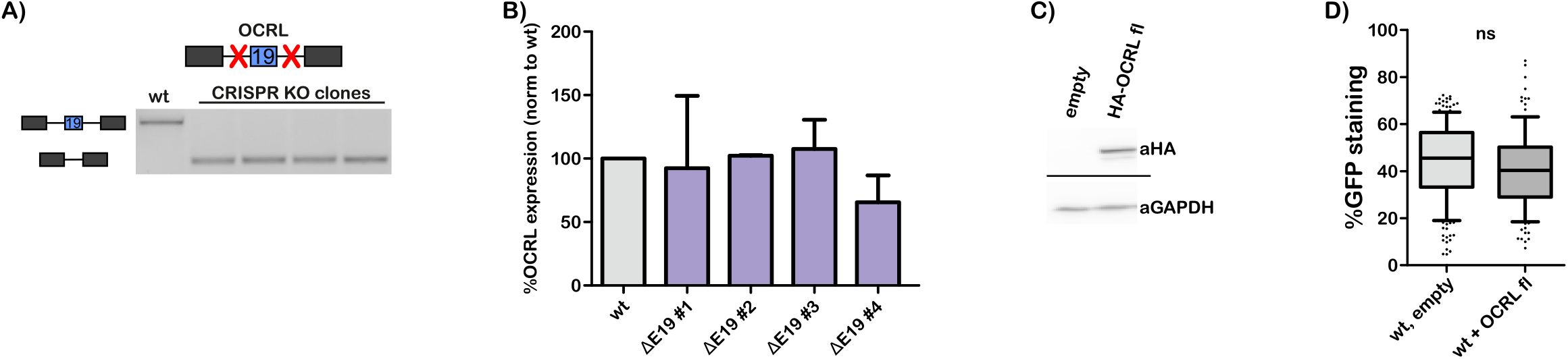
Characterization of OCRL KO clones and OCRL full length expression consequences. **A)** Genomic PCR validating KO of OCRL exon 19. **B)** Change in OCRL expression in the CRISPR clones analysed by RT-qPCR normalized to GAPDH and wt control. Shown are mean and SD of 2 independent RNAs from same cell line. **C)** WB showing OCRL full length construct expression. **D)** RUSH assay after expression of OCRL full length in wt cells. Whiskers show 10-90 percentiles, n>=100 cells, ns p>=0.05.

**Figure 4 - figure supplement 1:**
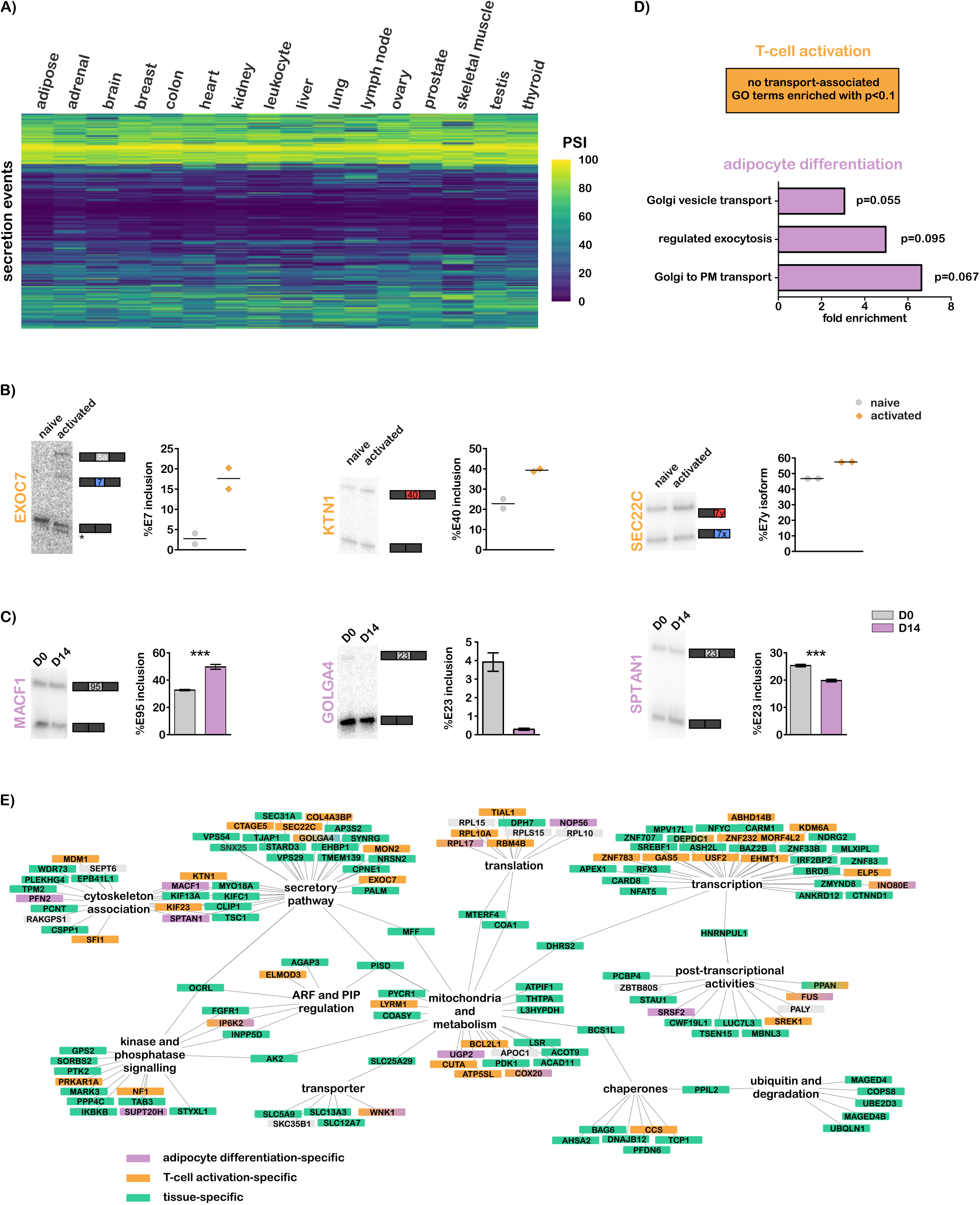
Further information regarding tissue‐ and differentiation-specific events. **A)** Tissue heatmap of RBP background events based on RNA-seq data from the lllumina Bodymap. PSI percent spliced in. **B, C)** RT-PCR validation of overlap events found differential in RNA-Seq data for T-cell activation (B) and SGBS adipocyte differentiation (C). T-cells: n=2, shown are replicates and mean. SGBS cells: Shown are mean and SD, n=6. **D)** Transport-associated biological process GO terms of the overlap events with all secretion events as statistical background. While T-cells show no particular enrichment, splicing events controlled during adipocyte differentiation are concentrated In post-Golgi processes. **E)** Network adapted from Fig. 1e, the genes with differentiation-specific events are marked In orange and violet for T-cell activation‐ and adipocyte differentiation-specific, respectively. Events that are differentially spliced between any two tissues are marked In green. If an event is present in multiple overlaps, a gradient fill is used.

